# Cortex-Wide Preservation of Multi-Stimulus Information across Degrees of Network Synchronization

**DOI:** 10.64898/2025.11.28.690948

**Authors:** Silviu-Vasile Bodea, Rodrigo Gonzalez Laiz, Rhȋannan H. Williams, Mariia Gladkova, Dominik Thalmeier, Bastian Rieck, Felix Sigmund, Marie Piraud, Afra Wohlschläger, Dominik Jüstel, Albrecht Stroh, Steffen Schneider, Gil Gregor Westmeyer

## Abstract

Sensory-evoked cortical responses vary with global network dynamics, yet the link between cortical state and stimulus processing remains unclear. Here, we introduce a paradigm that jointly decodes stimulus identity and network state from wide-field calcium imaging in mice undergoing multisensory and optogenetic stimulation across isoflurane-induced transitions from compartmentalized to synchronized activity. Effective dimensionality, a summary measure of network complexity, correlated well with anesthesia depth, while non-linear contrastive learning achieved >97% stimulus decoding accuracy across all states. Individual cortical regions maintained ≥82.5% accuracy even during deep anesthesia with prominent slow waves. Preservation of stimulus-specific information extended throughout the cortex, demonstrating that distinct representations remain decodable within synchronized networks. Direct optogenetic cortical stimulation exhibited state-invariant decoding performance, contrasting with anesthesia-dependent decline observed for sensory stimuli. Multi-stimulus cortical representations remain decodable across varying levels of network synchronization, with implications for brain-machine interfaces and clinical tools that assess preserved cortical responsiveness under variable arousal conditions.

---

Brain activity comprises continuously evolving electrical and chemical oscillations spanning a wide range of temporal and spatial scales^1^, with information processing essential for behavior superimposed on this dynamic background^2,3^.

During wakefulness, desynchronized spontaneous activity governs cortical networks, and compartmentalized cortical activity can be identified under lightly sedated conditions^4,5^. In contrast, synchronization of cortical networks and so-called slow waves (<1Hz) start to appear in quiet wakefulness^6^, growing in amplitude and low frequency power as cortical neurons become more synchronized in bursting activity during deep sleep^7,8^, and anesthesia^4,9–11^. Interestingly, slow-wave activity is also found in small cortical areas during wakefulness^12,13^.

The brain supports distinct patterns of neural activity organized into “network states” that correspond to different behavioral and cognitive demands across arousal levels, including anesthesia^2^. Here, we refer to “network states” as configurations of large-scale neural networks that support particular modes of information processing. Shifts in network state contribute to the heterogeneity of brain responses to identical stimuli, resulting in a broad spectrum of behavioral outcomes^14,15^. Studies in mice^12,16,17^ and humans^18^ demonstrate diminished responsiveness to sensory stimuli when cortical slow waves are prominent.

Cortical stimulations, such as transcranial magnetic stimulation (TMS), bypass sensory pathways to directly engage the intrinsic excitability of the cortex and its state-dependent dynamics^19,20^. In humans, TMS/EEG is used to modulate neuronal processes^21^, assess the natural frequency of oscillatory states^22^, and the cognitive impact of arousal^23,24^.

Most brain decoding approaches assume constant arousal or attentional levels when using machine learning to identify stimuli from neural activity^25–27^. Separate model architectures are typically employed for decoding arousal levels, including sleep/wake stages^28^ and anesthesia depth^29,30^.

Here, we combine pan-cortical calcium imaging with systematic variation of anesthesia depth and multisensory and optogenetic stimulation. We propose effective dimensionality (ED) as a straightforward measure of network complexity and employ a unified framework for joint stimulus and state decoding using CEBRA (Consistent EmBeddings of high-dimensional Recordings using Auxiliary Variables)^31^, which employs contrastive learning to generate low-dimensional latent embeddings conditioned on contextual variables. Our findings demonstrate that distinct multi-stimulus cortical representations persist across varying levels of network synchronization.

## Results

### Experimental setup

We built a custom wide-field fluorescence microscope to capture functional cortical activity at 20 frames/s from Camk2a-tTA;tetO-GCaMP6s transgenic mice (n=15; **Fig. 1a**). Calcium signals were obtained from GCaMP6s expression in excitatory cortical neurons under a CaMKII promoter^32^. Our intrasubject design alternated five-minute spontaneous activity recordings with five-minute stimulation sessions; for the latter, 50-millisecond stimuli were applied every 10 seconds (**Fig. 1b**, left and right, respectively; **Supplementary Movie 1**). All imaging data were registered to a 2D cortical mask derived from the Allen Mouse Brain Atlas (**Supplementary Fig. 1**).

**Fig. 1.**
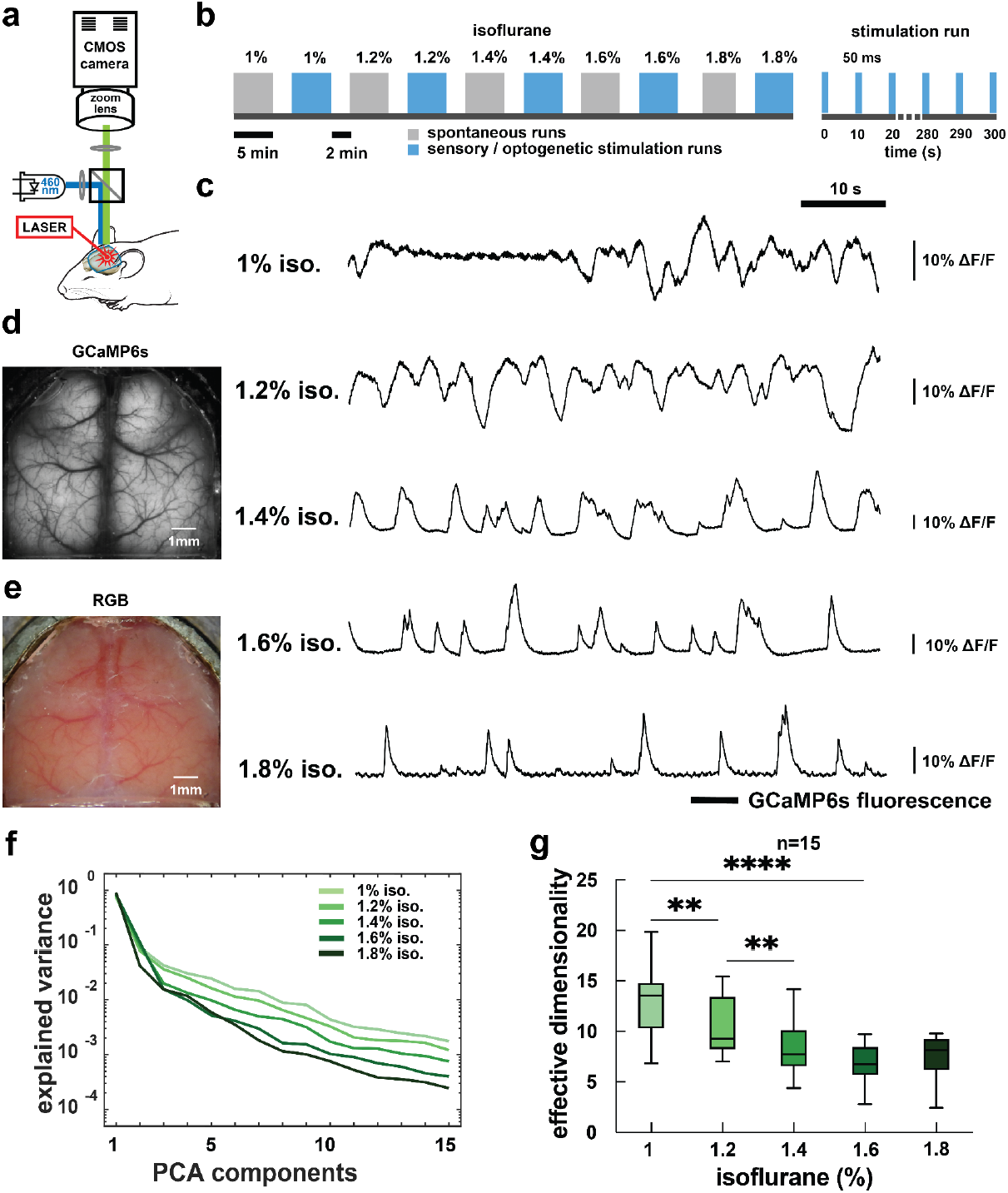
Experimental design and exemplary average activity traces representing spontaneous brain activity. **a**, Wide-field GCaMP6s fluorescence imaging setup. Imaging frames of 260×300 pixel resolution (∼35 μm/pixel) were acquired at 20 frames/sec. A 488 nm LED supplied light for GCaMP imaging. A fiber-coupled, 637 nm laser was used for optogenetic stimulation. **b**, Experimental design and stimulation schedule. 5-minute runs of spontaneous activity were followed by a 2-minute break and a 5-minute run during stimulation delivered in 50-ms pulses at 0.1 Hz using either natural stimuli (visual or somatosensory) or optogenetic stimulation in areas V1 or S1, respectively. One block of spontaneous /stimulation runs was repeated for isoflurane concentrations ranging from 1% to 1.8%, increasing in 0.2% increments. **c**, Average GCaMP6s cortical activity traces showing the progression from desynchronized activity to slow waves with increasing anesthesia. **d**, Representative epifluorescence image showing cortical signal acquired through a cranial window preparation. **e**, Corresponding RGB image, showing the view through the unopened dura **f**, Explained variance ratio per PCA component. Under high isoflurane anesthesia, fewer PCA components are required to explain the variance of the imaging dataset. **g**, A measure of effective dimensionality (see methods) was computed from spontaneous activity during the different depths of anesthesia. One-way ANOVA repeated measurements with Tukey’s multiple comparisons test for the comparisons between isoflurane concentrations (*, p<0.05, **, p<0.01, ***, p<0.001)

### Spontaneous activity dynamics

We first examined spontaneous network activity while incrementally raising the isoflurane (iso) concentration from 1% to 1.8%, in 0.2% steps (**Fig. 1b**). At 1% iso, average GCaMP6s traces revealed intricate dynamical patterns indicative of compartmentalized network activity **(Fig. 1c**, first and second rows). As the iso concentration increased, more wave-like patterns emerged (**Fig. 1c**, third row). During surgical-level anesthesia, exceeding 1.6% iso, slow waves became the dominant pattern (**Fig. 1c**, fourth and fifth rows, **Supplementary Movie 2 and Supplementary Fig. 6**).

### Effective dimensionality measures the complexity of spontaneous cortical activity

Principal component analysis (PCA) of the imaging data revealed that as anesthetic depth increased, fewer principal components were required to effectively represent the covariance structure of the data, reflecting increased cortical network synchronization (**Fig. 1f**). This finding led to the development of effective dimensionality (ED), a global complexity index for cortical activity. ED was derived from the Shannon entropy of the PCA eigenvalue distribution, normalized to a range between one and 50 PCA components (see methods). ED decreased significantly with increasing iso concentrations: from 13.1 at 1.0% iso to 10.5 at 1.2% iso (ANOVA and Tukey test for multiple comparisons, p=0.0013), and to 6.8 at 1.6% (p<0.0001). ED decreased further, plateauing between 1.6% and 1.8% iso (p=0.3; **Fig. 1g**). High ED values indicate greater compartmentalization of functional networks and more diverse cortical connectivity patterns, consistent with a lightly sedated state. As iso concentration increases to surgical levels, the decrease in ED reflects increased cortical synchronization and emergence of slow-wave activity (**Supplementary Movie 2**).

### Stimulus-specific cortical activation patterns

To examine how cortical network states affect sensory processing, we delivered white-light stimuli to the left eye (**Fig. 2a** and **Supplementary Movie 3**) and mild electrical pulses to the left hind paw for somatosensory stimulation^33–35^ (**Fig. 2d** and **Supplementary Movie 4)** following the schedule presented in **Fig. 1b**. At low anesthesia levels (1–1.2% iso), these stimuli consistently elicited responses in primary contralateral projection areas. Left visual stimulation activated the right V1 (primary visual) area (**Fig. 2b,c**; **Supplementary Fig. 2a**), and left paw stimulation activated the right S1 (primary somatosensory) hind limb region (**Fig. 2e,f**; **Supplementary Fig. 3a**). These activations were followed by ipsilateral responses in corresponding visual or somatosensory areas. At deeper anaesthesia levels (1.6–1.8% iso), stimulations initiated slow waves of calcium dynamics that spread across the entire cortex. Visual inputs evoked global responses originating from visual areas (**Fig. 2b,c**; **Supplementary Fig. 2b**), while somatosensory stimulations triggered global activations starting from central, somatosensory regions (**Fig. 2e,f**; **Supplementary Fig. 3b**).

**Fig. 2.**
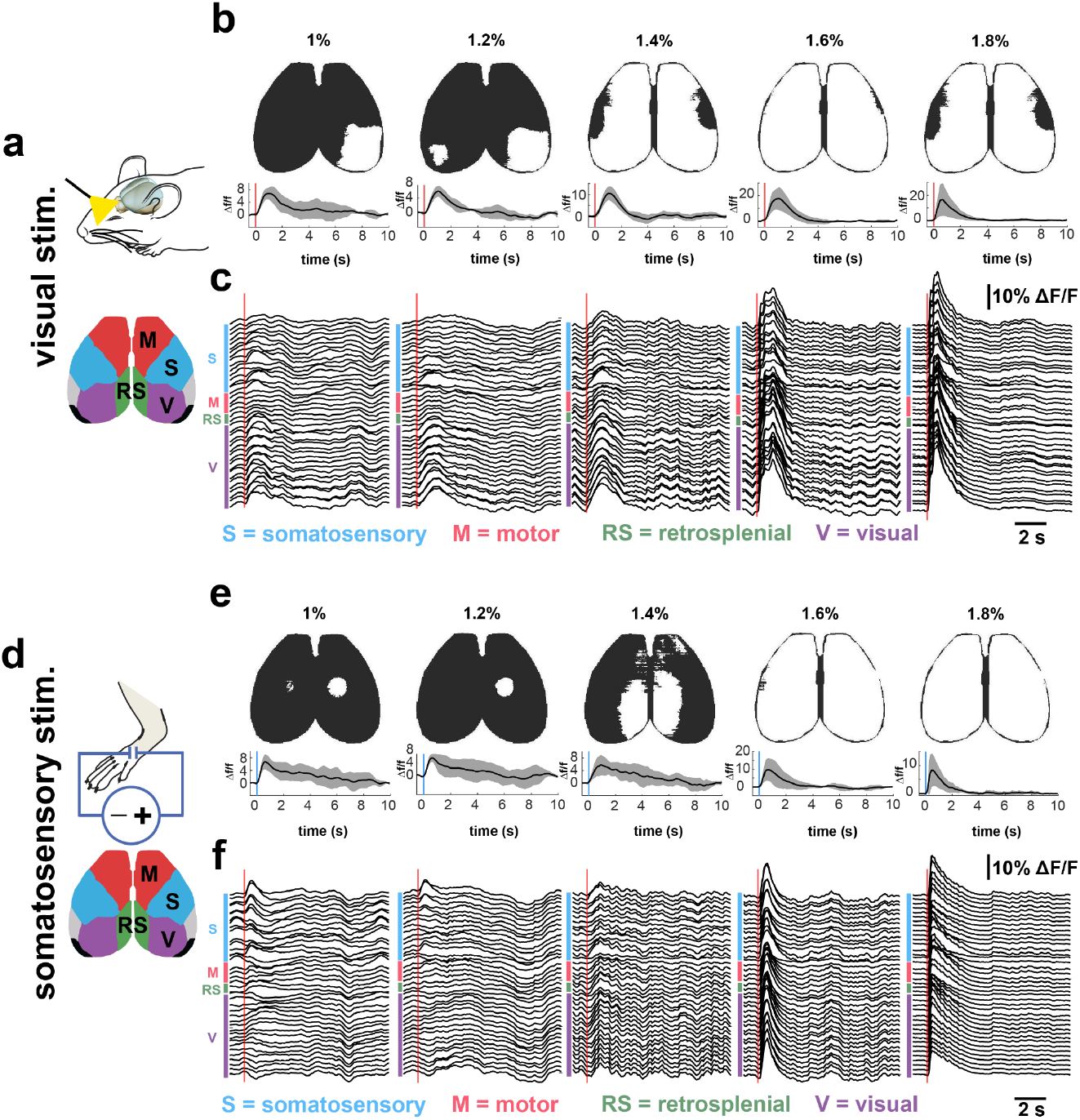
Modulation of stimulus-specific spatiotemporal brain activation patterns. **a**, Visual stimulation (*n*=5): 1 mW white light was delivered to the left eye in 50-ms-long pulses every 10 seconds, resulting in 30 stimulations per imaging run. Brain region segmentation map based on the Mouse Common Coordinate Framework (Allen Mouse Brain Atlas). V - primary visual cortex. **b**, Upper panel: binarized response 1 second after visual stimulation for increasing levels of isoflurane. The threshold was set at 50% of maximum activation in the V1 region under 1% iso. Lower panel: mean normalized signal change (Δf/f) in V1, with standard deviation (Stdev) as gray areas (*n*=5). **c**, Average signal change traces from sets of brain regions (S, somatosensory, M, motor, RS, retrospinal, V, visual) averaged over an entire run of a representative subject. From left to right: activation traces for increasing isoflurane anesthesia showing progression towards slow-wave activity. **d**, Somatosensory stimulations of the right paw (n=5): 1 mA, 50 ms long pulses were delivered every 10 seconds, resulting in 30 stimulations per imaging run. Brain region segmentation map based on the Mouse Common Coordinate Framework (Allen Mouse Brain Atlas). S - somatosensory cortex - hind limb. **e**, Upper panel: thresholded response 0.6s after hind paw stimulation (maximum intensity time index) for increasing levels of isoflurane. The threshold was set at 50% of maximum activation in the S1-HL region under 1% iso. Lower panel: Δf/f mean normalized signal change (Δf/f S1, with Stdev as gray areas (*n*=5). **f**, Average signal change traces from sets of brain regions (S, somatosensory, M, motor, RS, retrosplenial, V, visual) averaged over an entire run of a representative subject. From left to right: activation traces for increasing isoflurane anesthesia showing progression towards slow-wave activity.

**Fig. 3.**
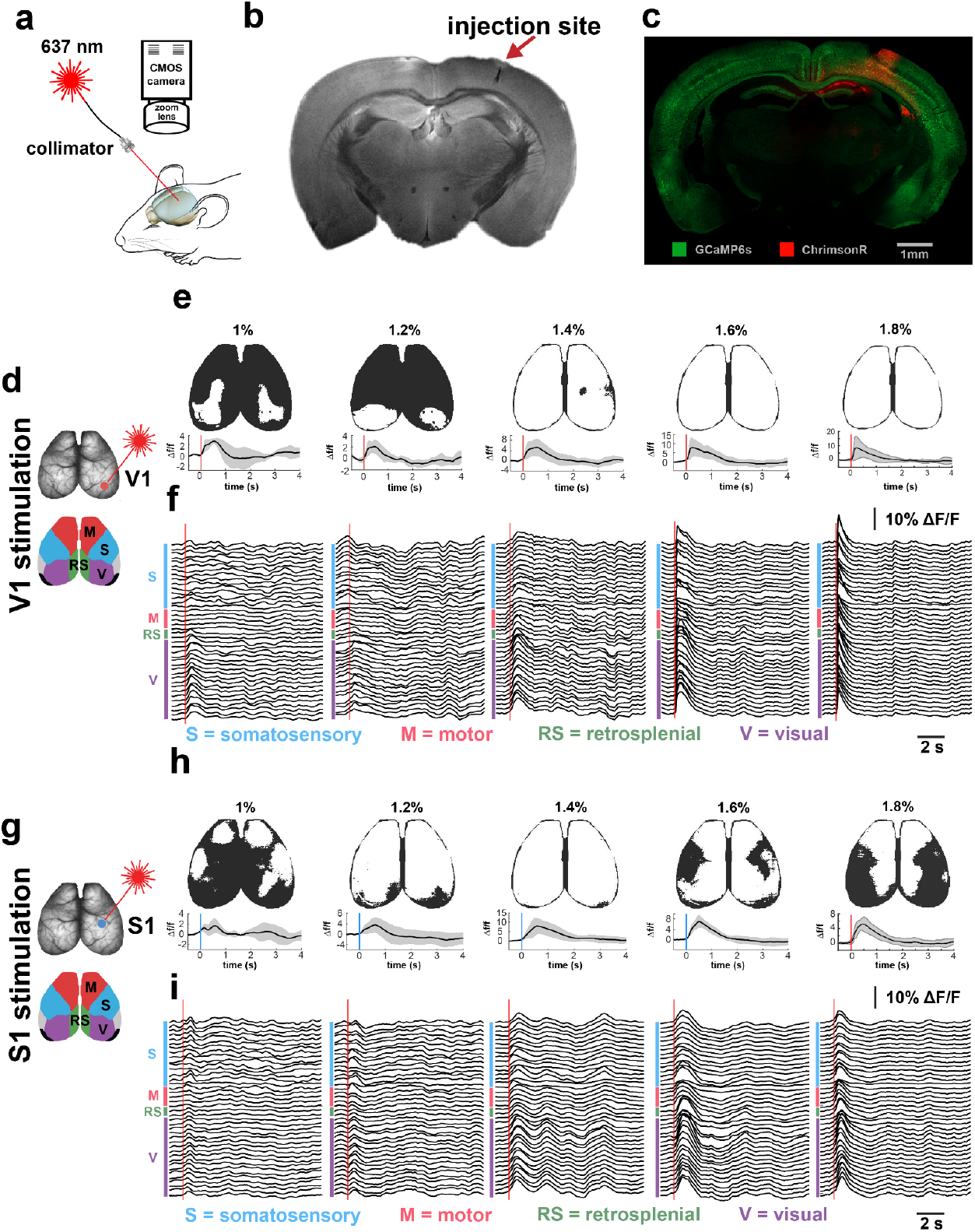
Activity patterns elicited by cortical optogenetic stimulation. **a**,**d**,**g** Schematic of the optogenetic stimulation of AAV-transduced Chrimson-R^Flag^ via a fiber-coupled 637 nm laser through an optical window in either V1 or S1. **b**, Coronal T_2_-weighted MRI tomogram showing the injection site in the somatosensory cortex of a representative mouse. **c**, Coronal section of a mouse brain immunostained with an anti-FLAG-Cy5 antibody against ChrimsonR^FLAG^ (red). Native GCaMP6s fluorescence is visible in green. **d**. Optogenetic stimulation site in the right visual cortex (red). 10 mW, 50-ms-long pulses were delivered every 10 seconds, resulting in 30 stimulations per imaging run. Brain region segmentation map based on the Mouse Common Coordinate Framework (Allen Mouse Brain Atlas). V1 - primary visual cortex. **e** Upper panel: binarized early response to V1 optogenetic stimulation for increasing levels of isoflurane. The threshold was set at 50% of maximum activation in the V1 region under 1% iso. Lower panel: mean normalized signal change (Δf/f) in V1, with Stdev as gray areas (*n*=5). **f**, Average signal change traces from sets of brain regions (S, somatosensory, M, motor, RS, retrosplenial, V, visual) averaged over an entire run of a representative subject. From left to right: activation traces for increasing isoflurane anesthesia showing progression towards slow-wave activity. **g**, Optogenetic stimulation site in the right somatosensory cortex (blue). Brain region segmentation map based on the Mouse Common Coordinate Framework (Allen Mouse Brain Atlas). S1-HL - primary visual cortex-hind limb. **h**, Upper panel: binarized early response to S1 optogenetic stimulation for increasing levels of isoflurane. The threshold was set at 50% of maximum activation in the S1-HL region under 1% iso. Lower panel: mean normalized signal change (Δf/f) in V1, with Stdev as gray areas (*n*=5). **i**, Average signal change traces from sets of brain regions (S, somatosensory, M, motor, RS, retrosplenial, V, visual) averaged over an entire run of a representative subject. From left to right: activation traces for increasing isoflurane anesthesia showing progression towards slow-wave activity.

### Optogenetic interrogation of state-dependent cortical dynamics

In a parallel set of experiments, we expressed the red-shifted opsin ChrimsonR^36^ in neurons of the right S1 and V1 cortices to modulate cortical dynamics (**Fig. 3a-c**). ChrimsonR stimulation (637 nm) triggers action potentials and calcium transients, increasing GCaMP6s signals from CaMKII-positive neurons. This enabled direct stimulation of excitatory networks in regions corresponding to the sensory projection areas under various network states (**Supplementary Movies 5 and 6**).

Under low anesthesia (1% and 1.2% iso), optogenetic activation patterns were more widespread than sensory-evoked responses, often involving associative areas (**Supplementary Figs. 4a, 5a**). Response amplitudes in V1 and S1 were lower compared with sensory stimulation (**Fig. 3e,h**).

**Fig. 4.**
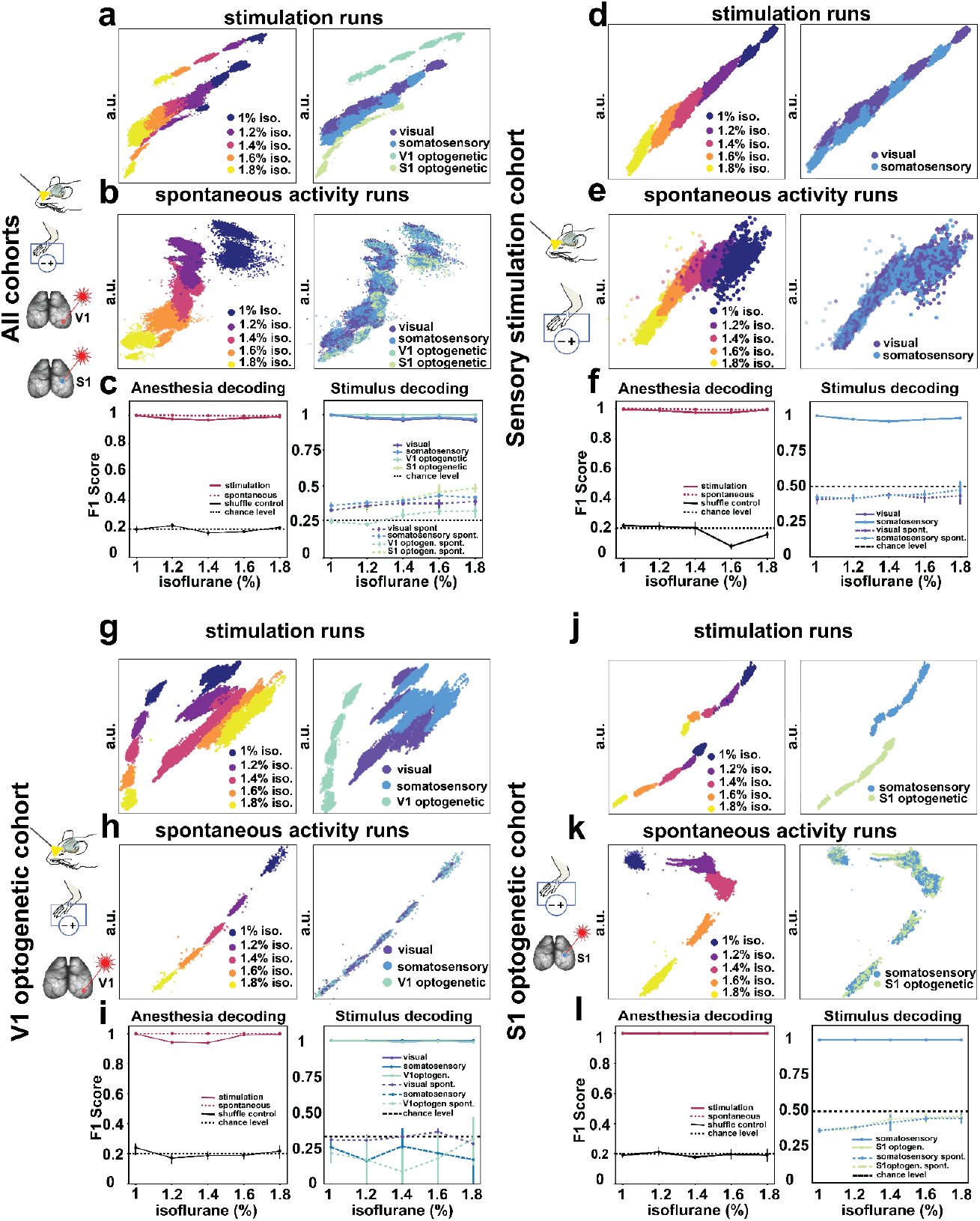
Non-linear embeddings across stimulus conditions and anesthesia levels. A 32-dimensional CEBRA-Discrete model was fitted across 20 discrete labels (4 stimuli x 5 anesthesia levels) using a 4-fold cross-validation. The first fold (arbitrary units, a.u.) is depicted here for all cohorts n=15 (**a-c**), the natural stimulation cohort n=5 **(d-f)**, the optogenetic V1 cohort n=3 (**g-i**), and the optogenetic S1 cohort n=3 (**j-l**). **a**, Joint embeddings of stimulation runs across all subjects, color-coded for anesthesia (left) and stimulation type (right). **b**, Embeddings from spontaneous activity runs of anesthesia (left) and stimulation type (right). **c**, Decoding performance (F1 score) for anesthesia decoding (left) and stimulus decoding (right) for stimulation runs (solid lines), and spontaneous activity runs (dotted lines). Anesthesia decoding was controlled with shuffled labels (black solid line). Broken lines indicate the results from the spontaneous activity control, the chance level with a black broken line. Stimulus decoding performance is not significantly influenced by increasing anesthesia (permutation test, p>0.1) **d**, Embeddings of stimulation runs, embeddings for anesthesia (left) and stimulation modality (right). **e**, Embeddings of spontaneous activity runs for anesthesia (left) and stimulation modality (right). **f**, Decoding performance for anesthesia decoding (left) and stimulus decoding (right). There was no significant difference in decoding performance across all iso. levels (permutation test p>0.1) **g**, Embeddings of stimulation runs, embeddings for anesthesia (left) and stimulation modality (right). **h**, Embeddings of spontaneous activity runs for anesthesia (left) and stimulation modality (right). **i**, Decoding performance for anesthesia decoding (left) and stimulus decoding (right). There was no significant difference in decoding performance across all iso. levels (permutation test p>0.1). **j**, Embeddings of stimulation runs, embeddings for anesthesia (left) and stimulation modality (right). **k**, Embeddings of spontaneous activity runs for anesthesia (left) and stimulation modality (right). **l**, Decoding performance for anesthesia decoding (left) and stimulus decoding (right). There was no significant difference in decoding performance across all iso. levels (permutation test p>0.1)

At higher iso concentrations (1.6% and 1.8%), optogenetic stimulation triggered propagating calcium waves that spread from the stimulation site across the entire cortex (**Supplementary Figs. 4b, 5b**). Optogenetically triggered waves propagated significantly faster than visually triggered waves (18.7±10.2 cm/s vs. 4.9±3.7 cm/s, p<0.0001; **Supplementary Fig. 6**), indicating distinct activation dynamics between optogenetic and sensory stimulation.

### Latent neural embeddings of anesthesia levels and stimulus-specific activations

We fitted a 32-dimensional CEBRA-Discrete model^31^ across 20 discrete non-ordered labels (4 stimuli x 5 anesthesia levels) with a four-fold cross-validation scheme, using data partitioned based on 10-second interstimulus intervals and 40 cortical regions (defined by the cortical mask in **Supplementary Fig. 1**). The resulting embedding spaces, depicted in **Figure 4**, illustrate how features for both anesthesia and stimulus decoding are represented in the same latent space. By treating anesthesia depth as an independent variable, we jointly conducted stimulus and network state decoding.

Joint embeddings for all 15 subjects (**Fig. 4a**) show high separability of the five anesthesia conditions for both stimulated activity (avg. F1 score 99.78%) and spontaneous activity recordings (avg. F1 score 98.15%). Stimulation did not impact anesthesia decoding (permutation paired F-test p>1) (**Fig. 4c left**). The anesthesia embedding spaces showed natural alignment with increasing anesthesia levels, achieving an R2 value of 98.61% (**Fig. 4a**, left). Anesthesia decoding scores remained consistently high across all five anesthesia levels (F-score 33, p>1) and significantly exceeded a shuffle control at the 20% chance level (avg. F1 score 19.74%; **Fig. 4c**, left). To test model robustness, we repeated the analysis on independent stimulation cohorts: sensory stimulation (**Fig. 4 d-f**), optogenetic V1 (**Fig. 4 g-i**), and optogenetic S1 (**Fig. 4 j-l**). Both stimulated imaging runs (F1 scores: sensory cohort 98.5%, optogenetic V1 cohort 97.3%, and optogenetic S1 cohort 99.9%) and spontaneous recordings (F1 scores: sensory cohort 99,6%, optogenetic V1 99.9%, and optogenetic S1 99.9%) showed high anesthesia decoding accuracy, significantly exceeding shuffle controls with F1 scores close to 20% (two-sample permutation test, p<0.0001).

Next, we investigated whether somatosensory, visual, optogenetic V1, and optogenetic S1 stimuli could be distinguished across anesthesia levels. The joint embedding space showed clear discrimination across all n=15 subjects (**Fig. 4a**, right) between sensory stimuli (F1 scores: visual 97.41%, somatosensory 98.11%) and between optogenetic perturbations (F1 scores: V1 99.99%, S1 99.99%). Classification performance was significantly above chance (permutation F-test, p<0.0001 for all stimulus modalities) and was not significantly influenced by anesthesia level (permutation F-test, p>1 for all modalities) (**Fig. 4c**, right).

We repeated this analysis in cohorts with randomized stimulus presentation. Differentiation was strong under light anesthesia, and stimuli remained distinguishable even under deep anesthesia despite traveling waves being the dominant pattern. In the *sensory stimulation* cohort (n=5; **Fig. 4d-f**), stimulus decoding performance was high for both visual (F1 score 97.75%) and somatosensory stimuli (F1 score 97.75%) (**Fig. 4f**, right), while spontaneous activity decoding remained at chance level (visual F1 score 42.08%, somatosensory F1 score 43.69%; **Fig. 4f**, right).

We investigated direct brain stimulation in two cohorts where V1 and S1 optogenetic stimulation were paired with visual and paw stimulation, respectively, testing whether optogenetic and sensory stimuli could be distinguished despite engaging the same cortical areas. In the *optogenetic V1* cohort (n=3; **Fig. 4g-i**), decoding was high for visual (F1 score 99.4%), somatosensory (F1 score 99.4%), and optogenetic V1 stimuli (F1 score 99.99%). Spontaneous activity decoding fell below the 33% chance level for all three stimuli (visual 31.6%, somatosensory 19.24%, optogenetic V1 21.16%; **Fig. 4i**, right). In the *optogenetic S1* cohort (n=3; **Fig. 4j-l**), stimulus decoding was high for both somatosensory (F1 score 100%) and optogenetic S1 stimulation (F1 score 100%), whereas spontaneous activity decoding fell below chance for both (somatosensory 41.43%, optogenetic S1 42.23%; **Fig. 4l**, right).

### Decoding features concentrate in ultralow frequencies

Having established that stimulus type and anesthesia depth can be decoded with high accuracy and minimal interaction at the global level, we examined which spatiotemporal features account for this decoding performance.

We first conducted a frequency-wise analysis (**Supplementary Fig. 7**) in which time traces were filtered into ultralow (0–0.5 Hz), delta (0.5–4 Hz), and theta (4–8 Hz) frequency bands.

High embedding similarity with raw data (**Supplementary Fig. 7a**,**b**) was primarily observed in the ultralow frequency band (**Supplementary Fig. 7c**,**d**), which also contributed most to anesthesia and stimulus decoding. This finding is consistent with the slow dynamics captured by GCaMP6s^37^.

In contrast, stimulus decoding accuracy was lower in the delta (**Supplementary Fig. 7e**,**f**) and theta (**Supplementary Fig. 7g**,**h**) bands. Anesthesia decoding in the theta band remained high for spontaneous data (F1 score >97%) but decreased to approximately 40% when stimulation was introduced (**Supplementary Fig. 7h**).

### Region-specific stimulus decoding

To investigate regional contributions to stimulus processing across anesthesia levels, we fitted an individual CEBRA model to each of the 40 anatomic brain regions (**Supplementary Fig 1**). Each region was subdivided in 40 equally spaced averaged subregions (**Supplementary Fig. 1**) across the three stimulation cohorts (**Figure 5**). This allowed us to use the same CEBRA architecture previously employed for the global cortical analysis.

**Fig. 5.**
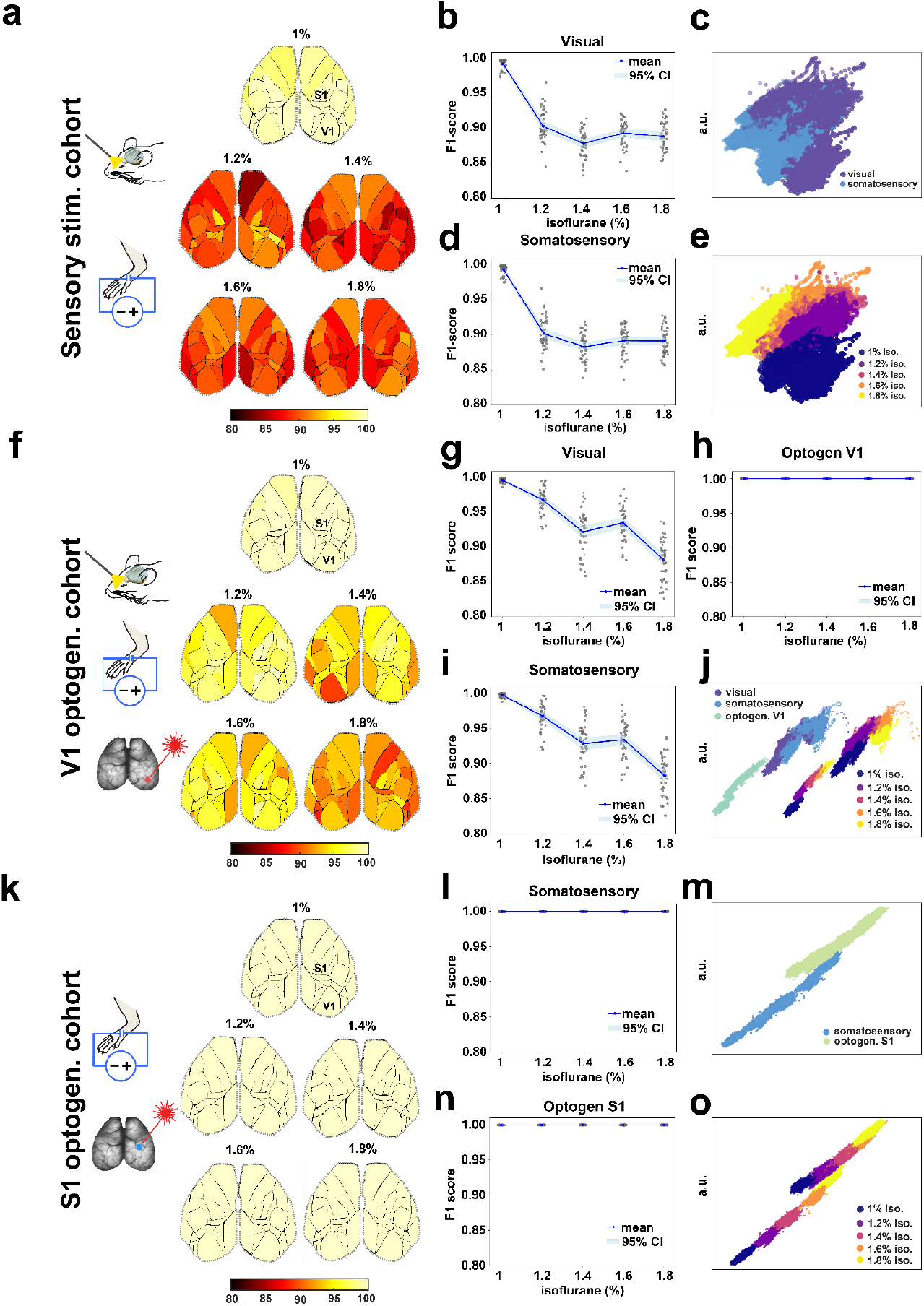
Cortical region-specific stimulus and anesthesia decoding. A CEBRA-Discrete model was fitted for each of the 40 regions of the Allen Mouse Brain Atlas. Models were trained with 4-fold cross-validation; the first fold is depicted here. Region-specific decoding was studied in randomized cohorts: the somatosensory cohort n=5 (**a-e**), the V1 optogenetic cohort n=3 (**f-j**), and the S1 optogenetic cohort n=3 (**k-o). a**, Sensory stimulation cohort cortical maps color-coded for decoding accuracy (F1 Score) across anesthesia levels progressing from compartmentalized activity to slow waves. Decoding accuracy decreases with anesthesia depth, with better performance in the primary S1-HL and V1 cortices. **b**, Decoding performance for visual stimuli dropped from 99.4% in 1% iso. to 90% in 1.2% iso. (permutation test, p<0.0001) and to 88.8% in 1.8% iso. (permutation test, p<0.0001). **c**, Embedding of stimulations. **d**, Decoding performance for somatosensory stimuli dropped from 99.4% in 1% iso. to 90.2% in 1.2% iso. (permutation test, p<0.0001) and to 89% in 1.8% iso. (permutation test, p<0.0001). **e**, Embedding of anesthesia. **f**, V1 optogenetic cohort cortical maps color-coded for decoding accuracy (F1 Score). Decoding accuracy decreases with anesthesia depth, with better performance in the primary S1-HL and V1 cortices. **g**, Decoding performance for visual stimuli dropped from 99.7% in 1% iso. to 96.9% in 1.2% iso. (permutation test, p<0.0001) and to 88.09% in 1.8% iso. (permutation test, p<0.0001). **h**, optogenetic V1 decoding accuracy was near-perfect and did not drop significantly, from 1% iso. to 1.8% iso. (p>0.01). **i**, Decoding performance for somatosensory stimuli dropped from 99.7% in 1% iso. to 96.7% in 1.2% iso. (permutation test, p<0.0001) and to 88.05% in 1.8% iso. (permutation test, p<0.0001). **j**, Stimulus and anesthesia embeddings for the V1 optogenetic cohort. **k**, S1 optogenetic cohort cortical maps color-coded for decoding accuracy (F1 Score) across anesthesia levels, showing near-perfect decoding accuracy for optogenetic S1 and somatosensory stimuli. **l**, Somatosensory decoding accuracy was near-perfect and did not drop significantly, from 1% iso. to 1.8% iso. (p>0.01). **m**, Stimulation embedding. **n**, optogenetic S1 decoding accuracy was near-perfect and did not drop significantly, from 1% iso. to 1.8% iso. (p>0.01). **o**, Anesthesia embedding showing high discrimination across somatosensory and S1 optogenetic stimulations.

Information for stimulus decoding was present not only in the primary visual (V1) and somatosensory (S1) projection cortices but in all 40 regions examined (**Fig. 6a, f, and k**). The lowest decoding performance was observed for somatosensory stimulation at 1.8% iso in the left latero-intermediate region, with an F1 score of 82.5%—still well above the 33.3% chance level (**Fig. 5i**, somatosensory). Unlike the global decoding results, discriminative performance in the sensory stimulation cohort declined with increasing anesthesia (**Fig. 5b and d**). Decoding accuracy decreased significantly for visual and somatosensory stimuli from an average F1 score of 99.4% at 1% iso to 90% at 1.2% iso (p<0.0001) and to 88.7% at 1.8% iso (p<0.0001). This finding was replicated in the optogenetic V1 cohort, which also included visual and somatosensory stimulation. Decoding accuracy declined by 3% between 1% and 1.2% iso (p<0.0001) and by 11.7% between 1% and 1.8% iso (p<0.0001) (**Fig. 5g, i**).

The decline in decoding performance correlated with a significant decrease in Euclidean distance between visual and somatosensory embedding space clusters (**Supplementary Fig. 8a, b**; 1% iso vs. 1.2% iso, p<0.0001). Additionally, the optogenetic V1 cluster was significantly closer to the visual than to the somatosensory cluster across most cortical regions (permutation test, p<0.05). At high anesthesia levels, this distance difference was no longer significant across large portions of the cortex (**Supplementary Fig. 9**).

In contrast, decoding performance for optogenetic stimulation did not decrease in the V1 optogenetic cohort (1% iso F1 99.9%; 1.8% iso F1 99.9%, p>0.99; **Fig. 5h**) or the S1 optogenetic cohort (1% iso F1 99.9%; 1.8% iso F1 99.9%, p>0.99; **Fig. 5l**). Euclidean distances between clusters increased slightly (**Supplementary Fig. 8b and c**), but this did not correspond to a significant increase in decoding accuracy.

## Discussion

Cortical stimulus representations undergo state-dependent modulation contingent upon arousal level, attentional allocation, and pharmacological perturbation of network dynamics. Here, we studied this interaction with a combined stimulus and state-decoding framework.

To systematically control network states, we employed isoflurane anesthesia to shift cortical activity from more localized non-synchronous patterns into widespread, slow-wave activity – a harmonized, coordinated oscillatory state. These anesthesia-induced states partially recapitulate natural arousal changes observed during drowsiness and sleep^38–40^, while offering greater experimental control than fluctuating behavioral states^41^.

Our imaging system captured GCaMP6s fluorescence from CaMKIIa-expressing neurons in superficial layers^32,42^ and ascending neuropil from L2/3 and L5^43^, linking microcircuit synchronization under anesthesia^10,44,45^ to cortex-wide stimulus processing through mesoscale imaging.

We developed a measure called effective dimensionality (ED), which is based on the entropy of PCA eigenvalues. ED can discriminate between isoflurane levels, indicating its potential for monitoring anesthesia depth. However, our experimental constraints prevented randomization of the isoflurane concentration sequence, and future validation would benefit from blood isoflurane concentration measurements^46^. ED alone could not distinguish between different stimulus types, indicating that identical dimensionality states support multiple sensory representations.

This limitation motivated exploration of non-linear computational approaches to extract stimulus-specific information across brain states. We leverage the CEBRA framework^31^ to learn embeddings of neural dynamics jointly conditioned on brain state and stimulus information. This approach differs from architectures that target sensory^25–27^ and network state decoding^28–30^ separately.

At the brain-wide level, the CEBRA-discrete model achieved >97% stimulus decoding accuracy across all anesthesia levels, including during slow-wave activity. The same embeddings enabled near-perfect prediction of anesthesia depth during both stimulated and spontaneous activity. Joint embeddings revealed distinct clustering patterns that remained separable even during maximal network synchronization. These findings suggest that the cortex maintains distinct stimulus representations even in highly synchronized, low-arousal states, consistent with previous studies^47–49^, though their relationship to predictions from studies emphasizing thalamic sensory gating^50–53^, and measurements under anesthesia finding preserved thalamocortical loops anchored in thalamic cores^54,55,56^, requires further investigation.

Region-wise analysis of models trained on each of the 40 single cortical regions showed a remarkably high decoding performance, also extending beyond primary sensory regions to heteromodal regions. This extends previous studies that found stimulus differentiation primarily within primary projection areas^48,49,57–59^.

In our experimental design, we included direct cortical stimulation via optogenetics to create an all-optical analog of TMS-EEG protocols, which probe cortical dynamics independent of peripheral sensory organs and ascending pathways^60^. This approach builds upon work combining optogenetics with optical hemodynamic readout^61^, and adds cell-type specificity by targeting CaMKII-expressing excitatory cortical neurons. Optogenetic stimulation maintained consistent decodability across anesthesia levels without the performance decline observed for sensory stimuli.

Notably, however, sensory stimuli also achieved high anesthesia decoding accuracy, and anesthesia depth could be classified accurately from spontaneous activity alone using CEBRA embeddings. These findings indicate that anesthesia depth tracking does not require direct cortical perturbation and can be achieved using simple global indices such as ED at the group level, or machine learning applied to recordings as short as 10 seconds. Furthermore, we found that decoding information was primarily represented in the ultralow frequency band (0–0.5 Hz), indicating that slow, large-scale cortical dynamics convey information about both sensory input and network state^37,62^.

These results suggest that similar approaches could be applied to human brain monitoring via EEG or fMRI, both during spontaneous activity and with simple sensory stimulation. This finding may also have practical implications for clinical monitoring, where high-density spatial or temporal sampling is constrained by hardware limitations and patient safety considerations. Models with joint brain state-stimulus embeddings may also enhance the reliability of brain-machine interfaces by maintaining decoding performance across fluctuating arousal states, a key consideration for integration of biological and technological circuits.

## Methods

### Mice

18 adult male B6 Camk2a-tTA;tetO-GCaMP6s^32^ mice (>8 weeks old, 25-40 g) were used in this study. Mice were genotyped for the GCaMP6s gene, and homozygous male mice were selected for experiments. GCaMP6s-positive animals were generously provided by the A. Stroh Lab (Univ. Mainz, Germany). 10 mice underwent viral injections for the expression of the optogenetic actuator ChrimsonR^FLAG^ (5 in the somatosensory cortex and 5 in the visual cortex), and glass window implantation was carried out 4 weeks after injection. 5 mice underwent glass window implantation only and received somatosensory and visual stimulation. Immunohistochemistry was done in 3 animals. Mice were housed under a 12-hour light/dark cycle. Food and water were provided ad libitum. All experimental procedures and animal husbandry conditions were approved by the Bavarian Government (Regierung Oberbayern).

### Surgical procedures

#### Viral injections

30 minutes before the start of the operation, mice were intraperitoneally administered 0.1 mg/kg buprenorphine (Temgesic 0.3 mg/ml, Eumedica). All surgical procedures were conducted under isoflurane anesthesia (3.5% for induction, 1.5-2% for maintenance). Anesthesia depth was regularly checked by corneal and toe pinch reflexes. Body temperature was maintained at 36.5°C with an electrical heating mat, and drying out of the cornea was prevented through the use of eye ointment (Bepanthen, Bayer). Mice were positioned in a stereotaxic device, and after hair removal, the skin was cleaned with Betadine (Braunoderm, Braun), and 50 μL lidocaine 2% (Braun) was injected subcutaneously for additional skin and periosteum anesthesia. A 7 mm incision was made, the fascia was gently pushed aside, and the skull was cleaned and allowed to dry for a few minutes. A dental drill was used to make a 400 μm in diameter burr hole while avoiding overheating or damage to the dura mater. A 33-gauge stainless steel injection cannula was lowered about 900 μm into the cortex, and 1 μL AAV solution was injected over a 5-minute period using a syringe pump (PHD 22/2000, Harvard Apparatus). The cannula remained in place for 10 minutes after the injection ended to allow the viruses to diffuse in the brain tissue before being slowly retracted. The incision was closed with the help of tissue glue (Vetbond, 3M), and lidocaine 2% was applied to the skin to prevent postoperative pain. 5 mg/kg Meloxicam (Metacam 2 mg/ml, Boehringer Ingelheim) was injected subcutaneously, and the animals were allowed to recover on heated mats until conscious. 5mg/kg Meloxicam was administered subcutaneously once a day for two days to provide postoperative analgesia.

### Glass window implantation

The preoperative procedures, analgesia, and anesthesia for the glass window implantation were carried out as described above. An 8 mm diameter oval skin flap was excised, and the fascia was pushed aside. A narrow stainless steel frame, slightly larger than the glass windows, was positioned and glued in place. Subsequently, a 10 mm skull window (Labmaker) was glued onto the skull with transparent UV-curing glue (Loctite 4305). The glue was slowly cured for 10 minutes by exposure to low-intensity 350 nm light. Subsequently, animals were allowed to recover, and postoperative analgesia was conducted as described above.

### Wide-field fluorescence imaging setup

The imaging instrument consisted of a variable zoom lens (Z16 APO, Leica) and a 0.5x objective (PlanAPO 0.5x, Leica 10447177) (**Figure 1a**). Imaging data was recorded at 20 frames/s and a resolution of 260×300 pixels with a sCMOS camera (Zyla 5.5, Andor). GCaMP6s excitation light was provided by a 3.3 Watt LED (UHP-T-LED-460, Prizmatix) bandpass filtered (ET 480/40 X, Chroma) and directed toward the brain with a long-pass dichroic mirror (T495lpxr-UF1, Chroma). GCaMP6s fluorescence was band-pass-filtered (ET 525/50 M, Chroma).

A Doric Lenses photometry console (D460-2002) was used to synchronize the camera and all stimulation devices. This console was responsible for triggering both the imaging frames and the stimulation.

### Imaging under Isoflurane

For anesthesia induction, mice were placed in a chamber that had been prefilled with 3.5% isoflurane. Once immobile, the animals were moved to a stereotaxic frame, and the isoflurane concentration was lowered to 1%. Eye ointment (Bepanthen, Bayer) was applied to prevent corneal drying. Animals were ventilated with a mixture of 80% air and 20% O_2,_ and body temperature was maintained between 36 and 37°C with a heating mat. After the animals stabilized in a sedated state (1% isoflurane), we acquired 5-minute spontaneous and stimulated imaging runs. Next, the anesthesia level was increased by 0.2%, and 2 minutes were allowed to pass for the higher isoflurane concentration to take effect. These steps were repeated for 5 isoflurane concentrations from 1% to 1.8% in 0.2% increments. Total imaging time was 50 minutes. After imaging, mice were kept on a heated mat until conscious.

### Stimulation schedule

We performed visual, hind paw, and optogenetic stimulations. All stimulations lasted 50 ms and were delivered every 10 seconds, resulting in 30 stimulations over a 5-minute imaging run (**Figure 1b**).

### Optogenetic stimulation

637 nm laser light for ChrimsonR stimulation was provided by a fiber-coupled LASER (S4FC637, Thorlabs). The laser light was guided through an optical fiber (P1-630Y-FC-2, Thorlabs) to a fiber optic collimator (F230FC-B-633, Thorlabs), which was manually positioned to achieve a 0.8 mm-diameter stimulation spot in the V1 or S1 regions. Typical laser intensity was 10-15 mW/mm^2^. Controls lacking expression of optogenetic channels were not included due to restrictions on animal numbers.

### Visual stimulation

Cool white light for visual stimulation was provided by a fiber-coupled LED (MCWF2, Thorlabs) and was guided through an optical fiber (M98L01, Thorlabs) to about 1cm in front of the left eye. The right eye was masked with light-opaque material.

### Hind paw stimulation

Mild electrical hind paw stimulation was done with a high voltage stimulus isolator (A365R, World Precision Instruments). The skin on the left hind paw was disinfected with 80% ethanol solution, and two 27-gauge platinum-iridium electrodes were implanted subcutaneously in the left hind paw. Stimulation intensity was set at 1 mA, and pulses lasted 50 ms. After the experiment, the paw was checked for possible minor hemorrhage, and the skin was again scrubbed with 80% ethanol.

### Immunohistochemistry

Perfused brains were incubated at 4°C for 24 hours in a 4% PFA solution and for 48 hours in a 30% sucrose solution before freezing at −80°C. 70 µm thick slices were prepared on a cryotome, and these were incubated for 1 hour at room temperature in a 1% BSA and 0.2% Triton X-100 in PBS solution for permeabilization/blocking. Slices were incubated at 4°C for 12 hours with an anti-FLAG Cy3 (A9594, Sigma Aldrich) monoclonal antibody (ChrimsonR was C-terminally tagged with a FLAG-epitope), which was diluted to a 1:500 ratio in 1% BSA and 0,2% TritonX-100 in PBS. Next, slices were washed 3 times for 5 min in PBS and incubated with DAPI solution, 10 mM diluted 1:1000 in 1% BSA and 0.2% Triton X-100 in PBS. After three 5-minute washes in PBS, the brain slices were mounted with Aqua Poly Mount (Polysciences, Warrington, PA). Cy5, DAPI, and GCaMP imaging were performed on an Axio Scan.Z1 (Zeiss) fluorescence microscope at 20x magnification.

### Image processing

Raw movies were motion-corrected using rigid transformations ^63^, and each imaging frame was registered to a cortical mask based on the Allen Mouse Brain Atlas ^64^ using manual control point selection and affine transformation implemented in a custom Matlab script.

### Effective Dimensionality (ED)

Effective dimensionality of a set of correlated variables refers to the equivalent number of orthogonal dimensions that would produce the same overall pattern of covariation ^65^. PCA component eigenvectors are orthogonal to each other and can be used to calculate effective dimensionality.

PCA allows decomposing a video into modes:

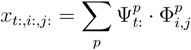

where 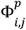 are the spatial modes, and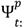 their temporal dynamics.

The total variance of the video averaged over time and space can be written as:

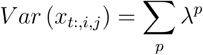

λ^*p*^, the eigenvalues of the PCA decomposition specify the contribution of each PCA component to the total variance. The magnitudes of the eigenvalues can be used to determine how many principal components are needed to adequately represent the imaging data. The first 50 PCA components were included in the calculation. To define the measure ED, we borrow from importance sampling, where the aim is to take weighted samples and determine how many effective unweighted samples these correspond to ^66,67^. The effective sample is a way to reduce a set of weighted elements to a set of unweighted elements. This concept lets us compute an effective number of dimensions from a set of weighted dimensions (principal components).

Analogous to the definition of effective sample size, we define ED as the entropy-based effective PCA dimensionality:

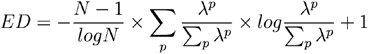

where *N* is the number of principal components. Defined in this way, ED is a normalized entropy measure bounded between 1 and N: the effective dimensionality equals N when all eigenvalues contribute equally and 1 when a single component explains all variance.

### Neural latent embeddings with CEBRA

#### Preprocessing

We reduce each imaging frame by clustering non-zero pixels (i.e., those inside the brain) into a fixed number of groups using k-means. The non-zero pixel coordinates are identified, clustered, and each pixel is assigned a cluster label. For every time frame, we then compute the mean activity across all pixels within each cluster, producing one representative signal per cluster. This procedure reduces the high-dimensional image to 1000 cluster-averaged signals while preserving the spatial organization of the cortex. For the multi-area experiments, we applied the same clustering procedure separately within each area (1–40). To ensure comparability and avoid size-related biases, we fixed the number of clusters to 40 per area, regardless of the region’s spatial extent. For all cohorts, we concatenate all data and apply min–max scaling to the range [–1, 1] to improve numerical stability during model training.

#### Cross-Validation

We used a 4-fold cross-validation scheme with non-overlapping groups of 200 frames, ensuring each group appeared exactly once in the test set across folds. No separate validation set was used. For control experiments, we shuffled the neural data in these 200-frame blocks per animal to disrupt the temporal alignment between neural activity and the labels (stimuli or anesthesia levels). Cross-validation was not performed in the region-wise experiments; instead, a simple train-test split was used on the same 200-frame blocks (75% train, 25% test).

#### Model training

We used the CEBRA framework (v0.4.0) to train the “offset10-model-mse” model, which is a 1D time-convolutional neural network consisting of an input convolution, a stack of residual CNN blocks with GELU activations, and a final convolution mapping to a 32-dimensional output. The network has a centered receptive field of 10 frames, enabling it to capture dependencies within a symmetric temporal window of the input sequence. The model was trained using the InfoNCE loss with a Euclidean distance metric for 30,000 steps, a batch size of 512, and a learning rate of 1e-4. We trained the CEBRA model using the discrete sampling mode, leveraging 20 unique labels (4 stimuli × 5 anesthesia levels) without imposing an order on these labels. After training, we extracted the embedding and classified each point using a logistic regression model implemented in scikit-learn (v1.7.2) with L2 regularization (C=1) on the same 20 labels, and reported both accuracy and F1 score after grouping the predictions either by stimulus or anesthesia level.

Embeddings were visualized by taking the first 3 dimensions of the learned 32-dimensional representation. Each point represents a window of 10 samples of neural activity embedded using the CEBRA encoder. The colors indicate stimulus identity or anesthesia level.

### Euclidean Distance Analysis

In **Supplementary Fig. 8**, we quantified representational separation by computing pairwise Euclidean distances between stimulus-class centroids in the embedding space for each brain region across three stimulation cohorts. For each region and anesthesia level, we computed the mean embedding (centroid) for each relevant stimulus class, calculated pairwise distances between centroids, and then averaged distances across regions. Statistics were computed with a nonparametric F-test and confidence intervals for the column means were computed using nonparametric bootstrap resampling (10,000 iterations), defined as the 5th and 95th percentiles of the bootstrap distribution.

In **Supplementary Fig. 9**, we computed pairwise Euclidean distances between class centroids for each brain region, animal, and anesthesia level. For each region and animal, we computed class centroids at five anesthesia levels, calculated all pairwise distances, and then averaged distances across animals and regions. Confidence intervals were obtained using the SEM across regions (±1.96 × SEM). The resulting curves show how centroid distances between stimulus pairs change with anesthesia, with shaded areas indicating 95% CIs.

To identify regions where V1 optogenetic stimulation diverged more from visual versus somatosensory stimulation, we ran permutation tests for each region and anesthesia level. Using the six relevant distance values (three animals × two distance types), we computed the observed difference between mean V1 optogenetic–somatosensory and V1 optogenetic–visual distances. We then enumerated all possible splits of the six values into two groups of three to form the null distribution. One-tailed p-values were defined as the proportion of permuted differences ≥ the observed difference. Regions with p ≤ 0.05 were displayed on brain-mask templates, with significant regions highlighted.

### Slow-wave speed determination

In 3 animals, visually (n=44) and optogenetically (n=30) initiated slow waves were recorded at a 100 frames/sec imaging rate to capture fast propagation patterns across the cortex. A custom brain mask (**Supplementary Fig. 6f**) was used to extract signals from 10 ROI located along a line from the occipital to the prefrontal cortex and positioned 0.8 mm apart. The ROI were binarized using a threshold that was set at 20% of the maximum fluorescence of each ROI, and the time to threshold was determined. The propagation speed was calculated by linearly regressing the time-to-threshold vs. distance traveled. The estimated speed was the slope of the regression line.

### Statistical analysis

Sample sizes were chosen using standards common in the field of imaging neuroscience for in vivo experiments. Statistics were reported using mean ± standard deviation. The statistical tests were used as indicated in the figure legends. Multiple comparisons with Tukey-Kramer tests (α=0.05) were carried out as follow-up tests after One-way and Two-way ANOVA, multiple comparisons were taken into account. Statistical tests were performed in Graphpad Prism 9.3.1 and using custom scripts in Python.

### F-test

A nonparametric permutation test was employed to assess differences in mean values across conditions. For the omnibus test, the observed test statistic (F-value) was computed from the original data. To generate the null distribution, condition labels were randomly permuted within the conditions (i.e., preserving the repeated-measures structure) 10,000 times, and the test statistic was recalculated for each permutation. The p-value was estimated as the proportion of permuted test statistics greater than or equal to the observed statistic. For post hoc pairwise comparisons, the same permutation procedure was applied to each pair of conditions, and p-values were corrected for multiple comparisons using the Holm–Bonferroni method. This approach makes no assumptions about the underlying data distribution and maintains the dependency structure inherent in the experimental design.

## Author contributions

S.B. conceptualized the study, acquired all data, performed data analysis, performed statistical analysis, interpreted results, drafted and edited the manuscript, and drafted and edited the figures; R.G.L. designed and implemented the CEBRA machine learning pipeline, performed statistical analysis, interpreted results, and edited the manuscript and figures; R.H.W., contributed to data acquisition, contributed to scientific content, and edited the manuscript and figures; M.G. advised on data analysis; D.T. developed the Effective Dimensionality measure; B.R. advised on data analysis; F.S. designed custom AAV for expressing Crimson Flag; M.P. advised on data analysis for Effective Dimensionality; A.W. contributed to the scientific content; D.J. advised on data analysis; A.S. provided transgenic animals, advised on study design, contributed to scientific content, and edited the manuscript. S.Sch. conceptualized and designed the CEBRA implementation, advised on data analysis, interpreted results, and wrote and edited the manuscript G.G.W. conceptualized the study, secured funding, supervised the project, provided scientific oversight and conceptual guidance, advised on data analysis, interpreted results, and wrote and edited the manuscript.

## Acknowledgements

We are grateful for the support of Deutsche Forschungsgemeinschaft (DFG), Project SPP 1665, DFG Project LF4D, the Helmholtz Association Initiative, Networking Fund on the HAICORE@KIT partition, and the European Research Council (ERC) under the European Union’s Horizon 2020 Research and Innovation Program Grant Agreement No. 715933.

## Data availability

The datasets generated during the current study are available from the corresponding author on request.

