## Supplementary Information for "Cortex-Wide Preservation of Multi-Stimulus Information across Degrees of Network Synchronization"

[1]Institute for Synthetic Biomedicine, Helmholtz Munich, 85764 Neuherberg, Germany
[2]Department of Bioscience, TUM School of Natural Sciences and TUM School of Medicine, Technical University of Munich, Munich, Germany
[3]Institute of Computational Biology, Helmholtz Munich
[4]Munich Center for Machine Learning (MCML)
[5]Institute of Neurogenomics (ING), Helmholtz Munich
[6]Technical University of Munich, TUM School of Computation, Information and Technology, Computer Vision Group
[7]Helmholtz AI, Helmholtz Munich
[8]Institute of AI for Health, Helmholtz Munich, 85764 Neuherberg, Germany
[9]Department of Diagnostic and Interventional Neuroradiology, Technical University of Munich
[10]Institute of Biological and Medical Imaging, Helmholtz Munich
[11]Institute of Computational Biology, Helmholtz Munich
[12]Chair of Biological Imaging at the Central Institute for Translational Cancer Research (TranslaTUM), School of Medicine, Technical University of Munich
[13]Institute for Physiology I, University Hospital Münster, University of Münster, Germany
[*]corresponding author

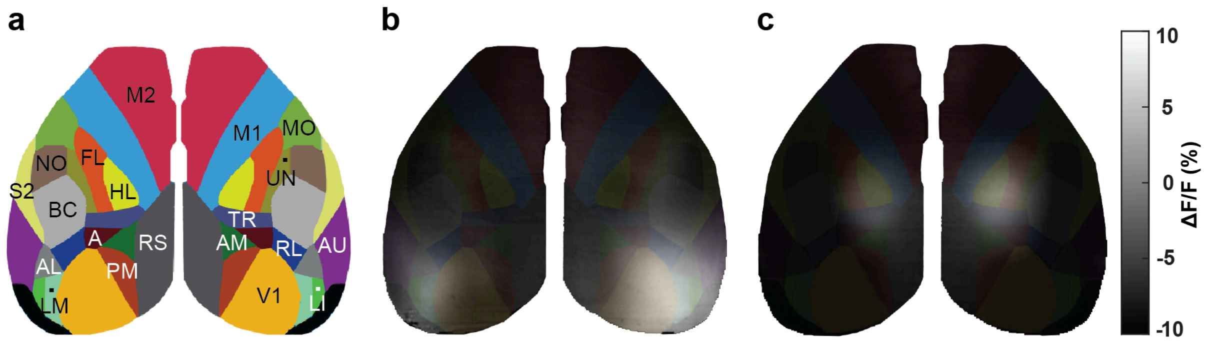


**Supplementary Fig. 1 | Coregistration of functional imaging data to a cortical mask based on the Allen Mouse Brain Atlas.**
**a,** Brain parcellation mask based on the Allen Mouse Brain Atlas with 20 brain regions per hemisphere. M1 - primary motor area, M2 - secondary motor area, HL - hind limb, FL - front limb, S2 - secondary somatosensory area, MO - mouth, NO - nose, TR - trunk, A - anterior association area, AL - anterolateral, AM - anteromedial, LM - lateromedial, LI - laterointermediate, PM - posteromedial, RL - rostrolateral regions of the extrastriate visual areas, AU - auditory, RS - retrosplenial, BC - barrel cortex, V1 - visual area, UN - somatosensory area unassigned. **b,** Average activation of 30 left eye visual stimulations in one representative subject. **c,** Average activation of 30 left paw somatosensory stimulations in one representative subject.

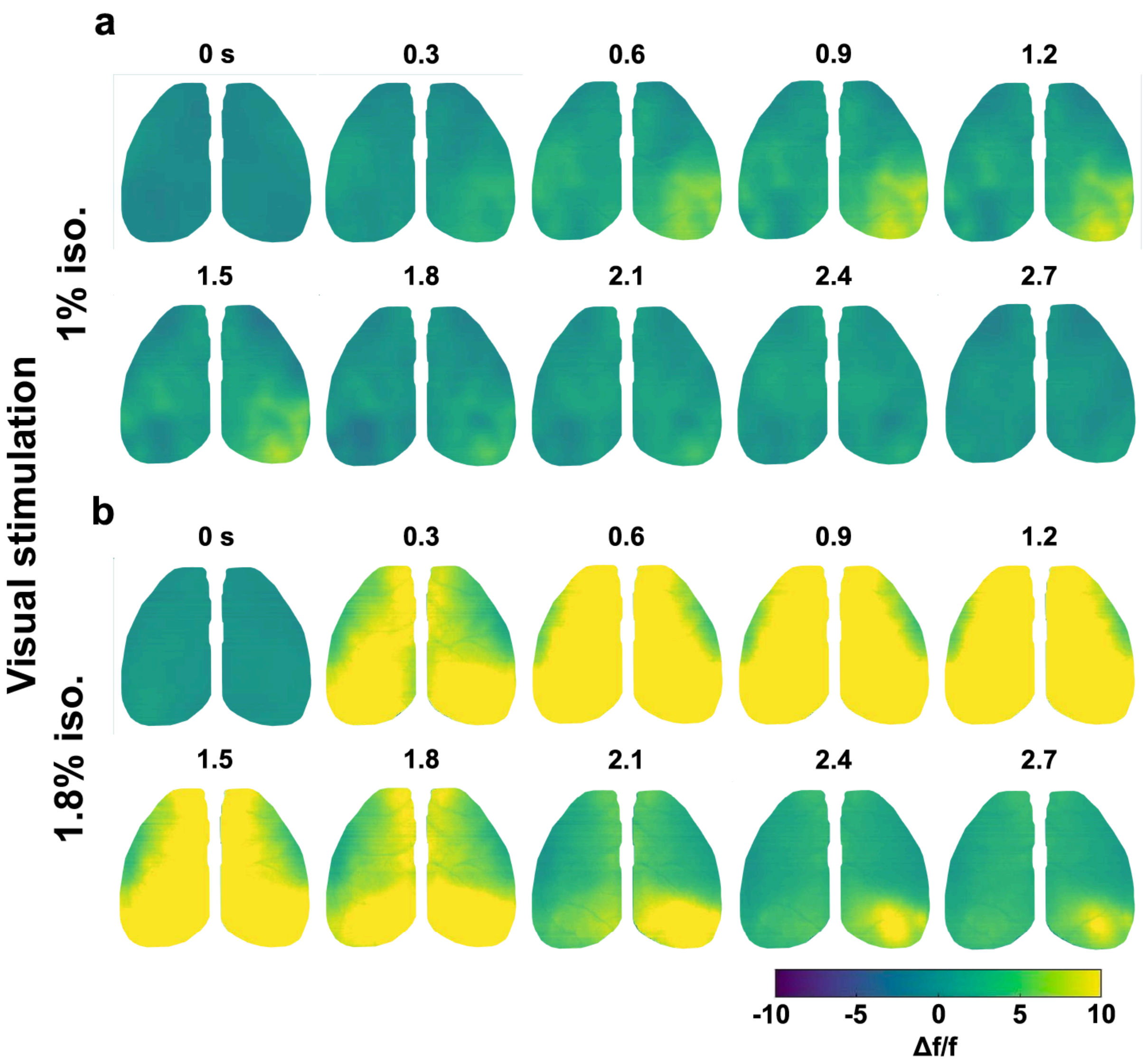


**Supplementary Fig. 2 | Responses to light stimulation of the left eye.**

**a,** Imaging time-series from an average of 30 visual stimulations in one representative subject during 1% iso. **b,** Imaging time-series from an average of 30 visual stimulations in one representative subject during 1.8% iso.

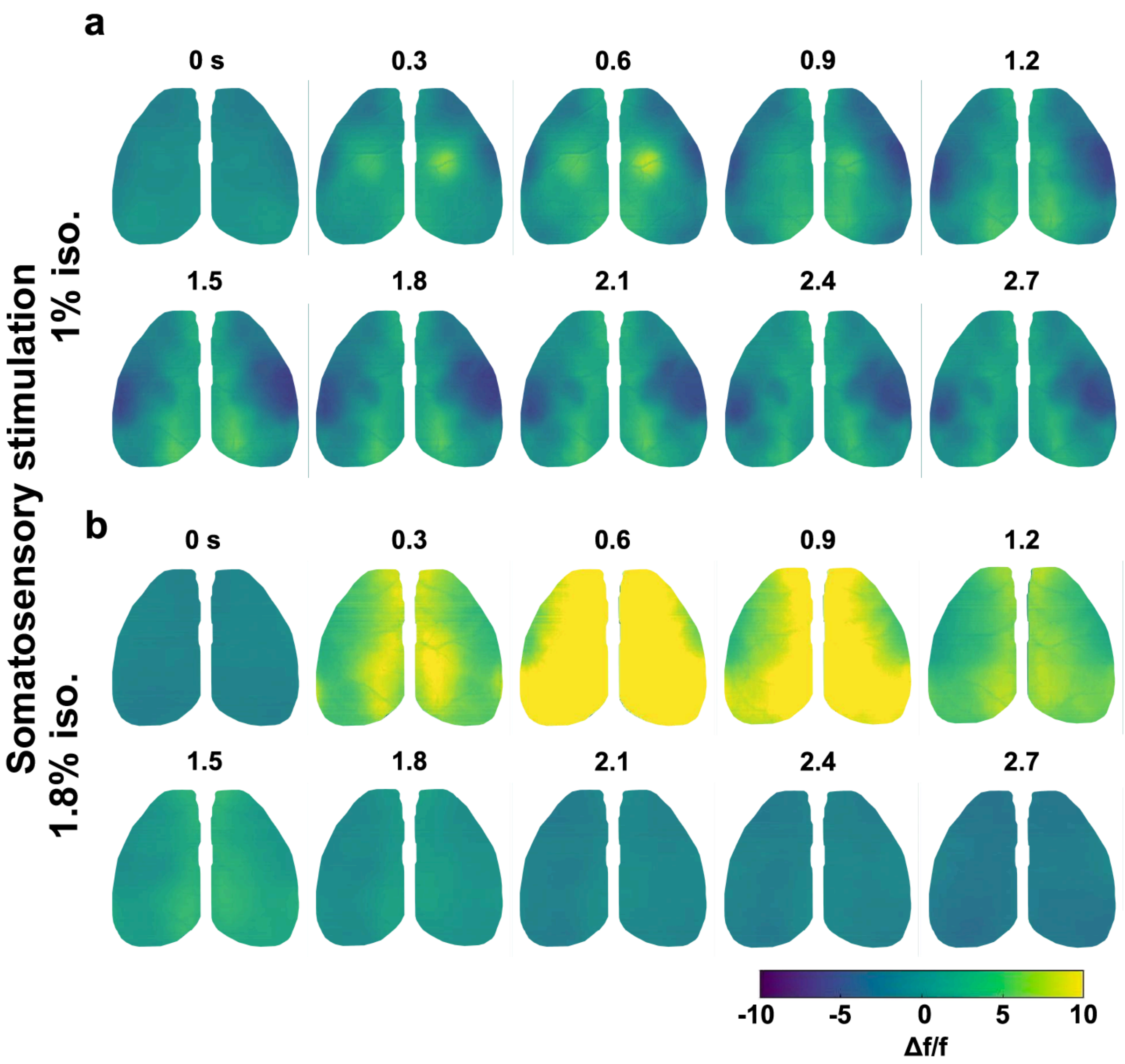


**Supplementary Fig. 3 | Responses to somatosensory stimulation of the left hind paw.**
**a,** Imaging time-series from an average of 30 hind paw stimulations in one representative subject during 1% iso. **b,** Imaging time-series from an average of 30 hind paw stimulations in one representative subject during 1.8% iso.

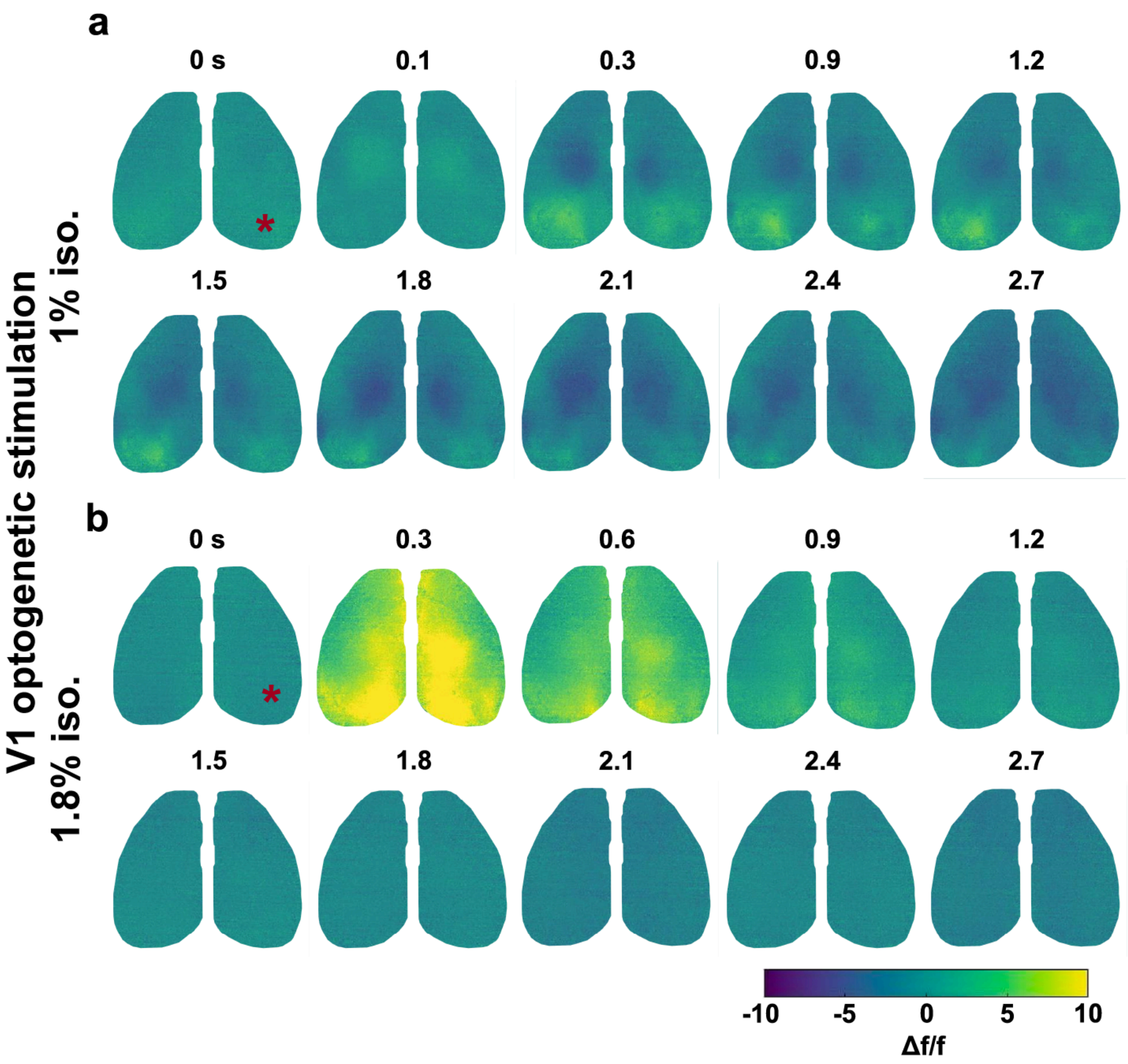


**Supplementary Fig. 4 | Cortical activation patterns after optogenetic stimulation of the right V1.**
**a,** Imaging time-series form an average of 30 optogenetic stimulations of the V1 region in one representative subject during 1% iso. **b,** Imaging time-series from an average of 30 optogenetic stimulations of the V1 region in one representative subject during 1.8% iso.

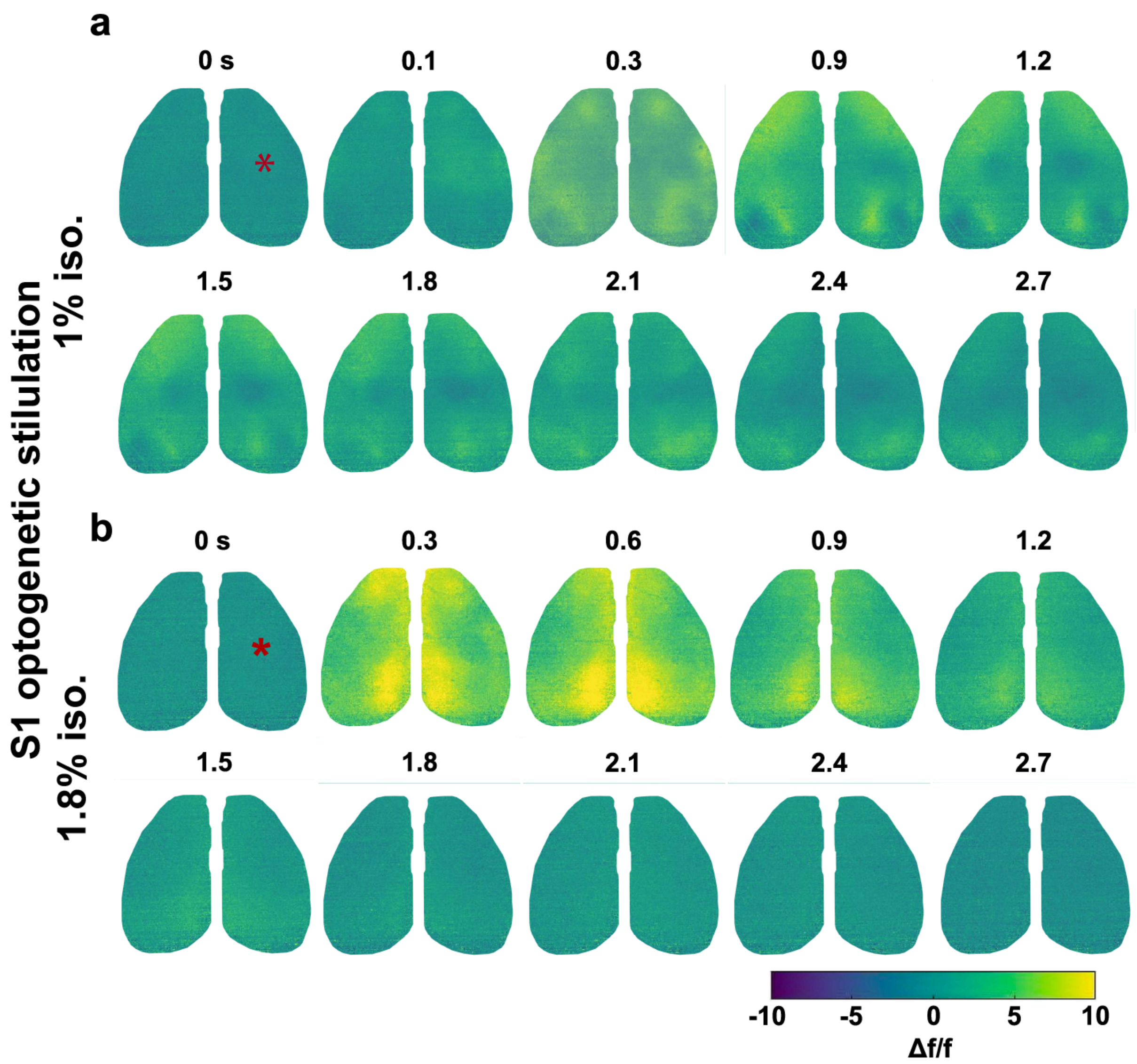


**Supplementary Fig. 5 | Cortical activation patterns after optogenetic stimulation of the right S1.**
**a,** Imaging time-series from an average of 30 S1 optogenetic stimulations in one representative subject during 1% iso. **b,** Imaging time-series from an average of 30 S1 optogenetic stimulations in one representative subject during 1.8% iso.

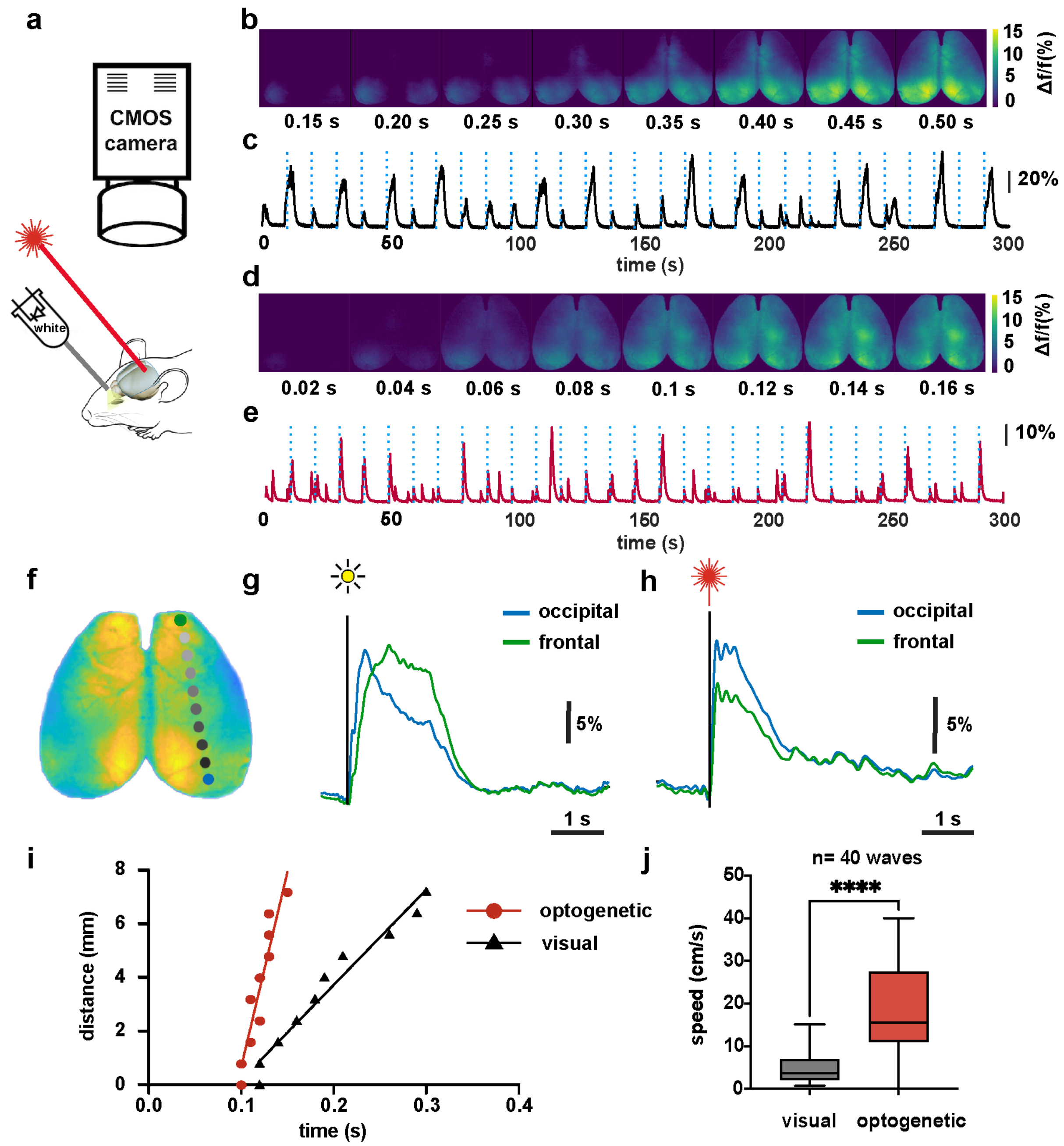


**Supplementary Fig. 6 | Optogenetic stimulation initiates traveling patterns of activation similar to slow waves.**

**a,** Responses to left visual white light and optogenetic right V1 stimulations were recorded at 100 frames/s in 3 mice under 1.6% isoflurane anesthesia. **b,** Montage showing cortical activation after visual stimulation. **c,** Average activation across the imaged cortex after visual stimulation (blue broken line). **d,** Montage showing cortical activation after V1 optogenetic stimulation. **e,** Average activation across the imaged cortex after V1 optogenetic stimulation (blue broken line). **f,** Brain mask with ROIs positioned from the occipital to frontal regions to assess propagation velocities. **g,** Averaged (n=44) occipital (blue) and frontal (green) time trace after visual stimulation. **h,** Averaged (n=30) occipital (blue) and frontal (green) time trace after optogenetic stimulation. **i,** Average (n=44) distance traveled was plotted against time for visually initiated (black, n=44) and optogenetically initiated (red, n=30) slow waves. The line represents the linear fit to the average data points. **j,** Propagation speed was determined for 44 visually initiated and 30 optogenetically initiated slow-waves and compared with an unpaired T-test (Welch's correction). The mean speed for visual waves was 4.9±3.7 and for optogenetic events was 18.7±10.2 cm/sec, $p<0.0001$.

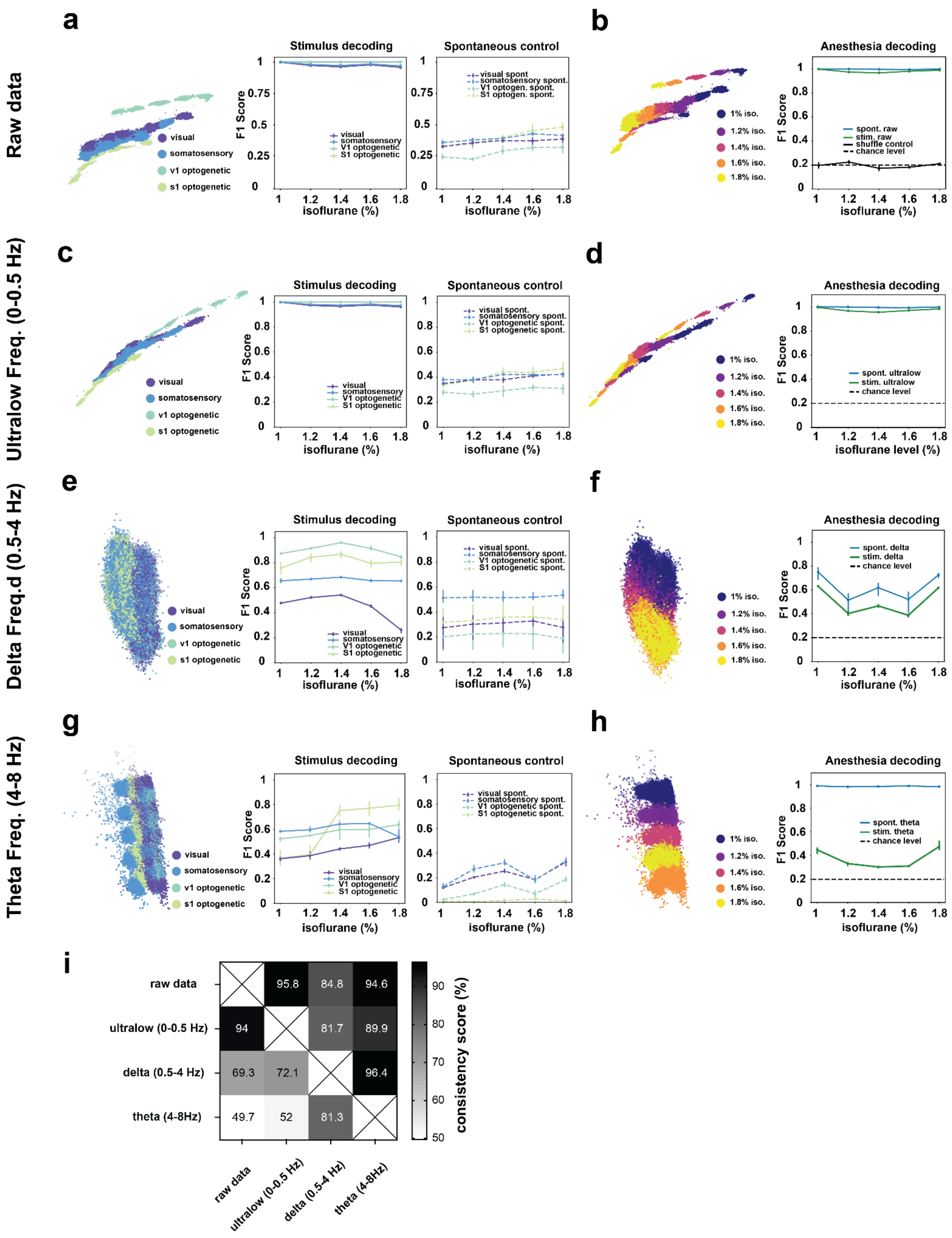


**Supplementary Fig. 7 | Frequency-dependent analysis.**

A CEBRA-Discrete model was fitted across 20 discrete labels (4 stimulus types x 5 anesthesia levels) using a 4-fold cross-validation. Stimulus and anesthesia decoding performance for raw data (**a-b**), ultralow (0-0.5 Hz) frequency band (**c-d**), delta (0.5-4 Hz) frequency band (**e-f**), theta (8-8Hz) frequency band (**g-h**). **a,** Raw data: left stimulus embedding, right stimulus decoding performance. **b**, Raw data: left anesthesia embedding, right anesthesia decoding performance. **c,** Ultralow frequency: left stimulus embedding, right stimulus decoding performance. **d**, Ultralow frequency: left anesthesia embedding, right anesthesia decoding performance. **e,** Delta frequency: left stimulus embedding, right stimulus decoding performance. **f**, Delta frequency: left anesthesia embedding, right anesthesia decoding performance. **g,** Theta frequency: right stimulus decoding performance. **h**, Theta frequency: left anesthesia embedding, right anesthesia decoding performance. **i,** Consistency score quantifying the similarity between the embeddings of the raw, ultralow, delta, and theta frequency datasets.

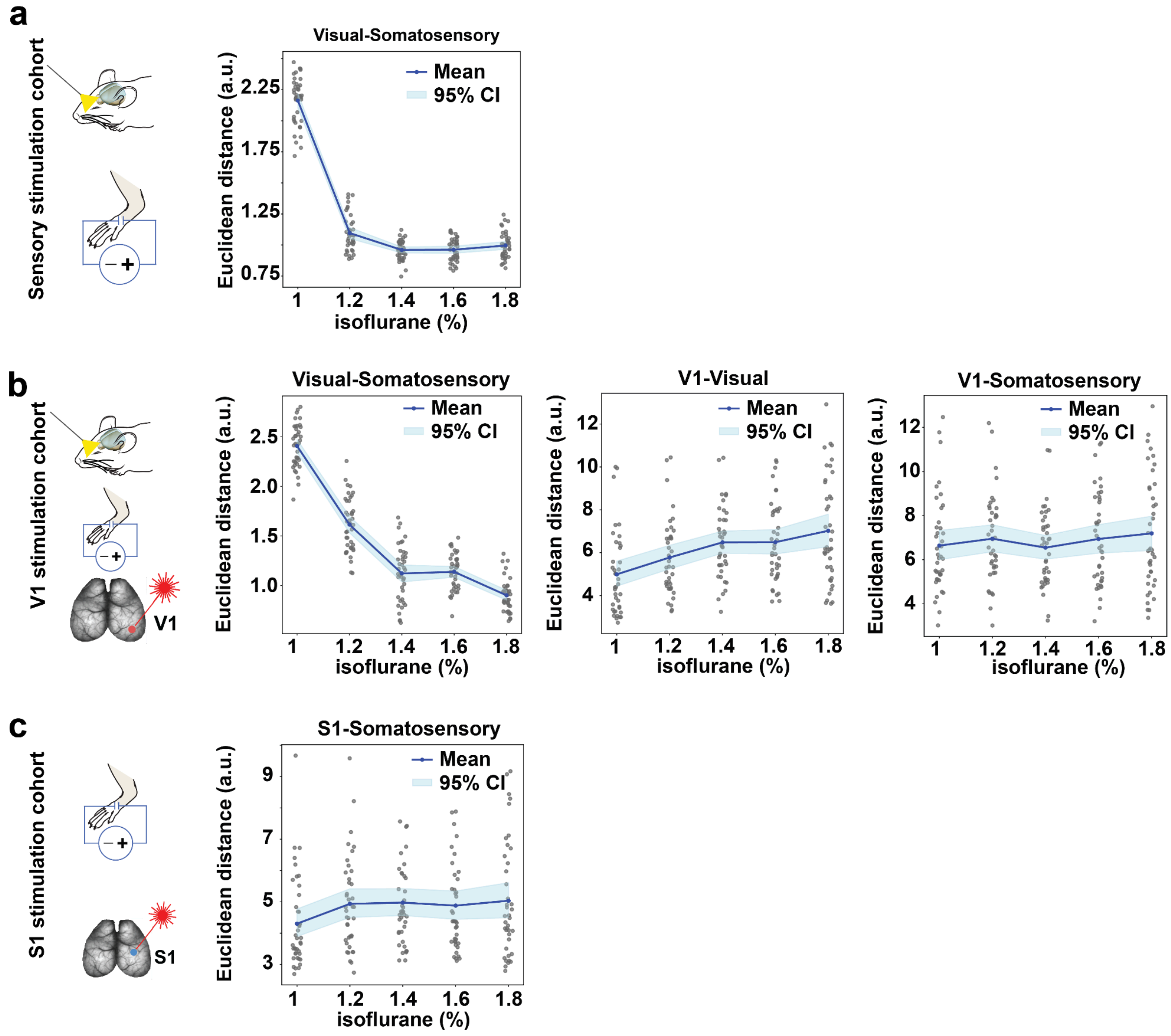


**Supplementary Fig. 8 | Euclidean distances. between stimulation clusters from region-specific embedding spaces.**
**a,** The Euclidean distance between visual and somatosensory clusters dropped from 2.16 (iso.1%) to 1.09 (iso. 1.2%) (permutation test p<0.0001). **b,** The Euclidean distance between visual and somatosensory clusters in the V1 optogenetic cohort dropped from 2.4 (iso.1%) to 1.6 (iso. 1.2%; permutation test p<0.0001) and to 0.9 (iso 1.8%; permutation test p<0.0001). **b,** The Euclidean distance between visual and somatosensory clusters in the V1 optogenetic cohort dropped from 2.4 (iso.1%) to 1.6 (iso. 1.2%; permutation test p<0.0001) and to 0.9 (iso 1.8%; permutation test p<0.0001). Distances between the V1 and somatosensory clusters did not increase significantly with increasing anesthesia. Distances between the V1 and visual clusters increased from 4.9 (iso. 1%) to 5.7 (iso. 1.2%; permutation test, p=0.001). **c,** The Euclidean distance between S1 and somatosensory clusters in the S1 optogenetic cohort increased from 4.3 (iso.1%) to 4.9 (iso. 1.2%; permutation test p<0.0001) and to 0.9 (1.8%; permutation test p<0.0001).

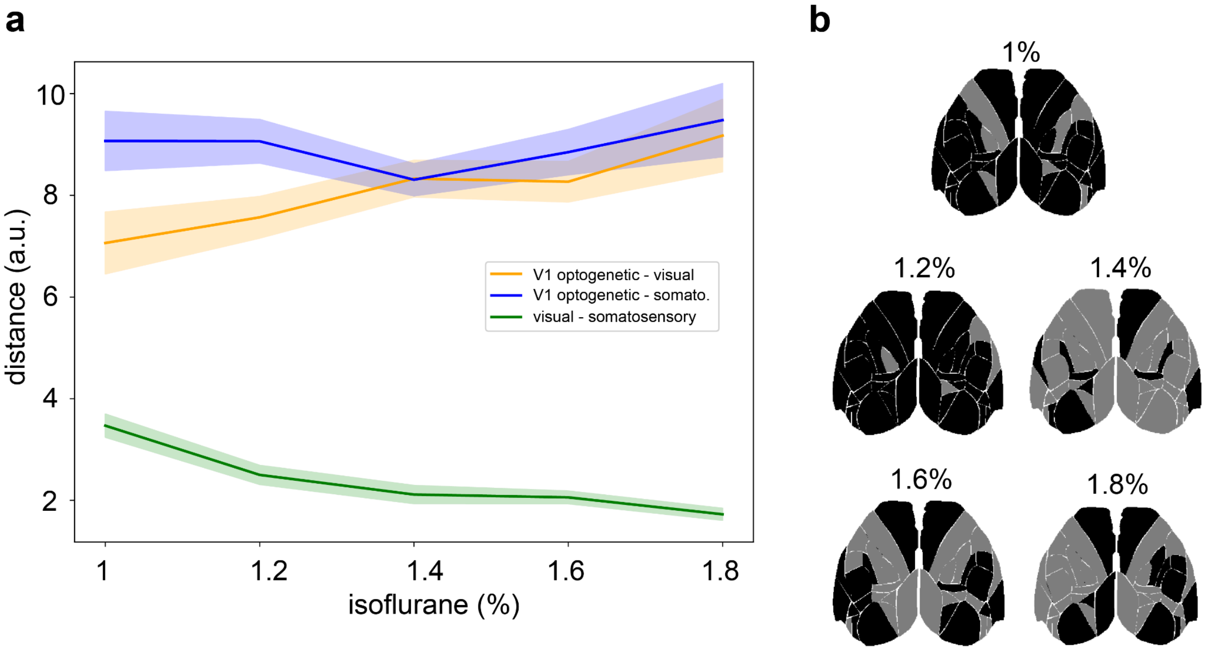


**Supplementary Fig. 9 | Distances between stimulus clusters in the embedding spaces of the V1 stimulation cohort.** We identified stimulus-dependent clusters in the CEBRA embedding spaces and measured their distances to characterize representational relationships. **a,** Euclidean distances between the stimulus-dependent clusters as indicated in the legend over the iso. levels. In 1% and 1.2% iso, the distances between the V1 and visual clusters (orange) are greater than those between the V1 and somatosensory clusters (blue). **b,** Distances were measured in the embedding spaces of all 40 CEBRA models. Cortical maps showing regions in black where the distances [V1 optogenetic - visual] (orange line in a) were significantly different from [V1 optogenetic - somatosensory] (blue line in a) (permutation test $p<0.05$). In gray areas, distances were not significantly different. In many regions, during 1% and 1.2% iso, V1 clusters are further apart from somatosensory clusters than visual clusters. As anesthesia increases, the distance difference becomes non-significant across large parts of the cortex. This could indicate that, during deeper anesthesia, representations remain stable and retain their differentiability.

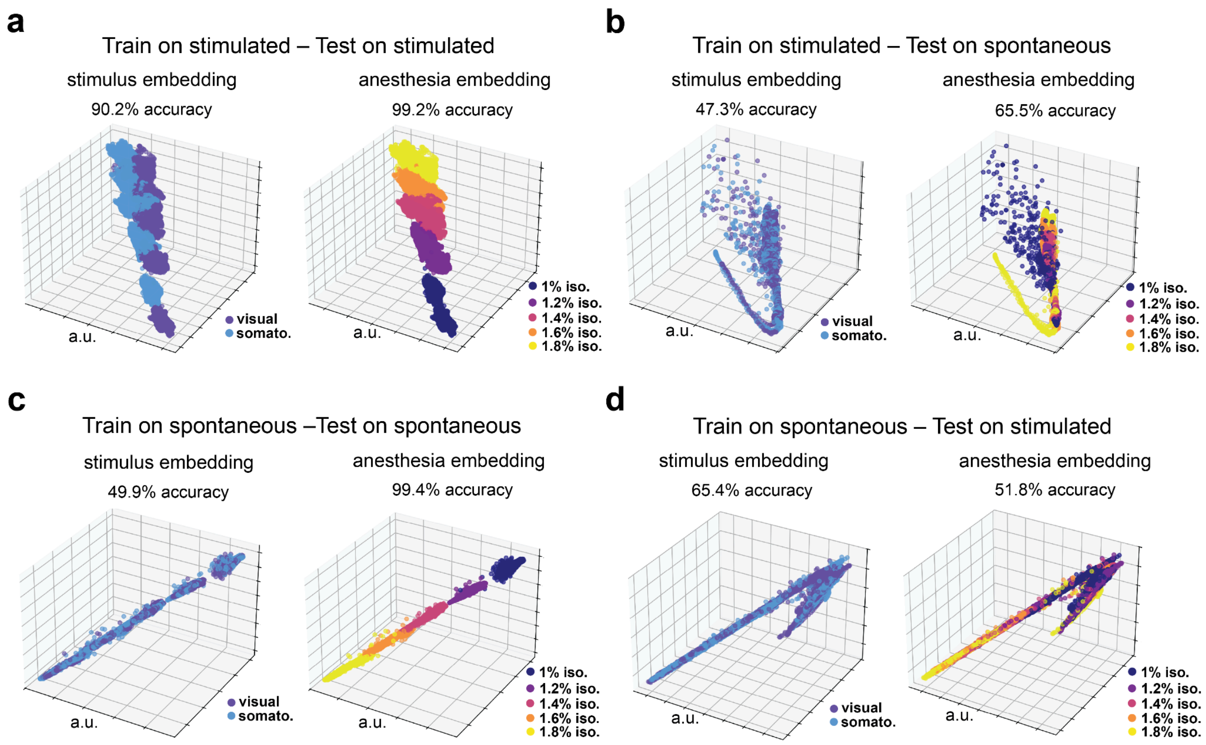


**Supplementary Fig. 10 | Cross-inference control for the sensory stimulation cohort.**
**a,** Stimulation and anesthesia embedddings for models trained on stimulated and evaluated on stimulated activity data. The decoding accuracy was 90.2% for stimuli (50% chance level) and 99.2% for anesthesia (20% chance level). **b,** Stimulation and anesthesia embeddings for models trained on stimulated and evaluated on spontaneous activity data. The decoding accuracy was 47.3% for stimuli (50% chance level) and 65.5% for anesthesia (20% chance level). **c,** Stimulation and anesthesia embeddings for models trained on spontaneous data and evaluated on spontaneous activity data. The decoding accuracy was 49.9% for stimuli (50% chance level) and 99.4% for anesthesia (20% chance level). **d,** Stimulation and anesthesia embeddings for models trained on spontaneous and evaluated on stimulated activity data. The decoding accuracy was 65.4% for stimuli (50% chance level) and 51.8% for anesthesia (20% chance level).
